# Towards fNIRS Hyperfeedback: A Feasibility Study on Real-Time Interbrain Synchrony

**DOI:** 10.1101/2023.12.11.570765

**Authors:** Kathrin Kostorz, Trinh Nguyen, Yafeng Pan, Filip Melinscak, David Steyrl, Yi Hu, Bettina Sorger, Stefanie Hoehl, Frank Scharnowski

## Abstract

Social interaction is of fundamental importance to humans. Prior research has highlighted the link between interbrain synchrony and positive outcomes in human social interaction.

Neurofeedback is an established method to train one’s brain activity and might offer a possibility to increase interbrain synchrony. Consequently, it would be advantageous to determine the feasibility of creating a neurofeedback system for enhancing interbrain synchrony to benefit human interaction.

In this study, we investigated whether the most widely employed metric for interbrain synchrony, namely wavelet transform coherence, can be assessed accurately in near real-time using functional near-infrared spectroscopy (fNIRS), which is recognized for its mobility and ecological suitability for interactive research. To this end, we have undertaken a comprehensive approach encompassing simulations and a re-evaluation of two human-interaction datasets. Our findings indicate the potential for a stable near real-time measurement of wavelet transform coherence for integration durations of about one minute. This would align well with the methodology of an intermittent neurofeedback procedure.

Our investigation lays the technical foundation for an fNIRS-based system to measure interbrain synchrony in near real-time. This advancement is crucial for the future development of a neurofeedback training system tailored to enhance interbrain synchrony to potentially benefit human interaction.

## 1. Introduction

Social interaction is vital not only for our survival but also for our physical and mental health (Cohen, 2004). Our interactions range from verbal to nonverbal communication, and they promote cooperating with each other, bonding with each other, as well as learning from one another. Impaired social interaction abilities, on the other hand, are detrimental to us and have been associated with, e.g., learning disorders (Kavale & Forness, 1996) or psychiatric disorders (Schilbach, 2016).

While traditionally, social neuroscience research has focused on single subjects reacting to social stimuli, in recent years, the focus has shifted towards investigating subjects in direct interaction (e.g., Hari & Kujala, 2009; Dumas, 2011; Schilbach et al., 2013). This so-called “second-person neuroscience” provides not only increased ecological validity, but has also revealed distinct neural signatures associated with direct interactions (Redcay & Schilbach, 2019).

During interaction, we perceive each other’s behavior, react to each other, and adapt our behavior accordingly. Our behavior is the means through which information about our mental states is conveyed to the other agents we are interacting with (Kingsbury & Hong, 2020). Importantly, these mental states are associated with our brain states. The dynamic relationship between our behaviors and percepts thereof is therefore reflected in our brains: the neural signals of each interacting agent become coupled and aligned through the agents’ reciprocal behavior (Dumas, 2011; Hasson et al., 2012; Hasson & Frith, 2016).

To investigate how two brain signals are coupled during social interaction, interbrain synchrony (IBS) is typically determined (e.g., Czeszumski et al., 2022). IBS signifies the temporal co-occurrence and alignment of neuronal signals, thus indicating that the signals are temporally related. IBS is thought to emerge both from a dynamical systems theory perspective as well as from a perspective of predictive coding and active inference: Dynamical systems theory indicates that when two sufficiently similar dynamical systems are coupled, they become synchronized (Friston & Frith, 2015a). Additionally, using the framework of active inference, it has been suggested that two brains predict each other during interaction and synchronization emerges when two predictive coding schemes are coupled (Friston & Frith, 2015b; Koban et al., 2017). According to this framework, IBS facilitates mutual prediction by minimizing the social prediction error (Friston & Frith, 2015b; Hamilton, 2020; Hoehl et al., 2020; Kingsbury et al., 2019; Koban et al., 2017; Pan et al., 2023; Shamay-Tsoory et al., 2019).

By now, IBS has been examined across a wide range of interacting social agents, including both humans (see Czeszumski et al. (2022) and Lotter et al. (2023) for recent meta-analyses of IBS and cooperative behavior) and non-human animals (bats, W. Zhang & Yartsev, 2019, and mice, Kingsbury et al., 2019).

To measure IBS during an interaction, it is necessary to image two brains simultaneously, i.e., to perform so-called hyperscanning (Montague et al., 2002; see Czeszumski et al., 2020 for an overview). Hyperscanning has by now been established in all major neuroimaging technologies, including functional near-infrared spectroscopy (fNIRS) (e.g., Babiloni & Astolfi, 2014). A major advantage of fNIRS for hyperscanning is that it is mobile and can be applied in nearly all natural environments. Further, participants are able to sit, stand, or move in their natural upright position while undergoing fNIRS neuroimaging. Two or more individuals can face each other directly and talk or interact manually with each other in a naturalistic way. While this is also possible with electroencephalography (EEG), fNIRS has the added advantage of increased spatial resolution combined with considerably lower susceptibility to motion artifacts compared to EEG (Kohl et al., 2020; Pinti et al., 2020).

Using fNIRS, IBS has been tested in a range of hyperscanning studies investigating different aspects of social interaction, both in adults as well as in adult-child interaction (e.g., Nguyen et al., 2020; Reindl et al., 2018), mainly in dyadic settings, but also in larger groups, e.g., 3-on- 3 person competition (J. Yang et al., 2020). The types of social interactions investigated with fNIRS and IBS include cooperative tapping experiments (e.g., Cui et al., 2012; Funane et al., 2011), teaching (e.g., Nozawa et al., 2019; Pan et al., 2018, 2020; Zheng et al., 2018), creative problem solving (e.g., Xue et al., 2018), counseling (Y. Zhang et al., 2018), and speed dating (Yuan et al., 2022); see also Czeszumski et al. (2022) for a recent meta-analysis. Specifically, the strength of IBS as measured by fNIRS has been associated with problem-solving success (e.g., Cui et al., 2012; Nguyen et al., 2020; Pan et al., 2017; Reindl et al., 2018) and prosociality during cooperation (Hu et al., 2017), and with in-group bonding during competition with another group (J. Yang et al., 2020). Finally, in the realm of teaching, IBS has been correlated with learning outcomes in teaching settings (e.g. Pan et al., 2018, 2020; Zhang et al., 2022). Taken together, these studies suggest that IBS is associated with interactive performance, both with the information transfer between individuals during interactions, as well as with associated social motivations.

Given these reported positive associations between IBS and human interaction, it is reasonable to hypothesize that enhancing IBS may lead to valuable improvements in human interaction. This may be especially the case where human interaction is impaired, as is the case in, e.g., certain psychiatric conditions (e.g., Schilbach, 2016).

One option to enhance a neural state is training it with the help of neurofeedback. Neurofeedback is an established method, which provides participants with information about their neural state with the goal of aiding them in regulating it (e.g., Sitaram et al., 2017). It is established in EEG and fMRI and has also recently gained traction in the domain of fNIRS (Kohl et al., 2020; Soekadar et al., 2021; Godet et al., 2023; Kohl et al., 2023). During a neurofeedback session, a participant’s neural state is measured in near real-time, and information about this state is then fed back to the participant in the form of, e.g., a visual thermometer or an auditory tone. With the help of feedback, participants can learn to self-regulate certain aspects (e.g., activation level, connectivity) of their brain activity. Learning self-regulation through neurofeedback training has been demonstrated to change the behavior associated with the trained brain signal. This intervention has also been shown to improve clinical symptoms in psychiatric and neurological patients (e.g., Sitaram et al., 2017).

Neurofeedback allows participants to gain access to their brain state. In the case of two interacting participants, this would mean receiving additional information about both participants’ brain states or the direct relation of both brain states, as would be the case in IBS neurofeedback. Normally, during an interaction, one person’s brain state can only be communicated to another person behaviorally, i.e., through speech or actions (e.g., Dumas, 2011; Kingsbury & Hong, 2020). By adding neurofeedback to an interaction, the natural information transfer between two brains through behavior can be enhanced with information about the interacting brain states.

Given the potential benefits of providing neurofeedback of IBS, it has been proposed and called for recently several times (Gvirts Provolovski & Perlmutter, 2021; Holroyd, 2022; Liu et al., 2019; Moreau & Dumas, 2021; Saul et al., 2022; Vanutelli & Lucchiari, 2022). However, it has only been sparsely attempted so far. Using EEG, pioneering work has been done in artistic contexts, where participants were visually fed back information about their brain state during interaction (Dikker et al., 2019; Dikker et al., 2021; Kovacevic et al., 2015). Recently, Müller et al. (2021) have demonstrated controllability over theta band synchrony in EEG, and an interface for EEG hyperfeedback has been presented (Chen et al., 2021). In animals, a proof-of-principle study demonstrated control of IBS in pigeons using invasive electrophysiological recordings (L. Yang et al., 2020). Using fNIRS in humans, the concept of cross-brain neurofeedback has been described (Duan et al., 2013); however, in this proof-of-principle study, two subjects were instructed to activate their brain activity higher than their partner, which is different from controlling IBS directly.

Although the scientific and possible clinical benefits of IBS neurofeedback are accepted, there is as yet no working implementation in the domain of fNIRS. This is likely due to methodological challenges, including modest signal quality, small effect sizes, and the ongoing refinement of real-time signal processing, which has yet to reach standardization (Kohl et al., 2020).

Here, we aim to take the first step in constructing an fNIRS-neurofeedback setup by testing a near real-time hyperfeedback analysis pipeline. We aim to demonstrate that the to-be-fed-back IBS signal, which carries the neural information, can be determined in near real-time. This means that the IBS metric needs to be reasonably calculable in short time intervals. Additionally, it needs to be robust enough when measured in short intervals to carry meaningful IBS information on the single-dyad level. We test these prerequisites of IBS neurofeedback by simulating and re-analyzing dyadic fNIRS data in near real-time and assessing the feasibility of such a setup.

## 2. Methods

### 2.1 Wavelet Transform Coherence analysis

In fNIRS experiments, wavelet transform coherence (WTC) is by far the most used metric for investigating interbrain synchrony (Czeszumski et al., 2022; Hakim et al., 2023). For calculating the coherence in a pseudo-real-time fashion, meaning calculating it as it would be necessary in a real-time situation, we adapted the currently used WTC approaches. Wavelet transform coherence was estimated using the “wcoherence” function from the Matlab wavelet toolbox, which follows the work of Grinsted et al. (2004). We used the default setting of Morlet/Gabor wavelets and averaged over a period band of interest. For all analyses in this manuscript, we have chosen a period band of interest of 6s to 14s, which corresponds to the upper and lower period boundaries utilized in the original analyses of the re-analyzed studies of this manuscript (see Fig. 1 and section 2.4). We have chosen 14 “VoicesPerOctave” (14 subbands/wavelets) instead of the default 12 to achieve the best coverage of the 6-14s period band of interest.

**Fig. 1:**
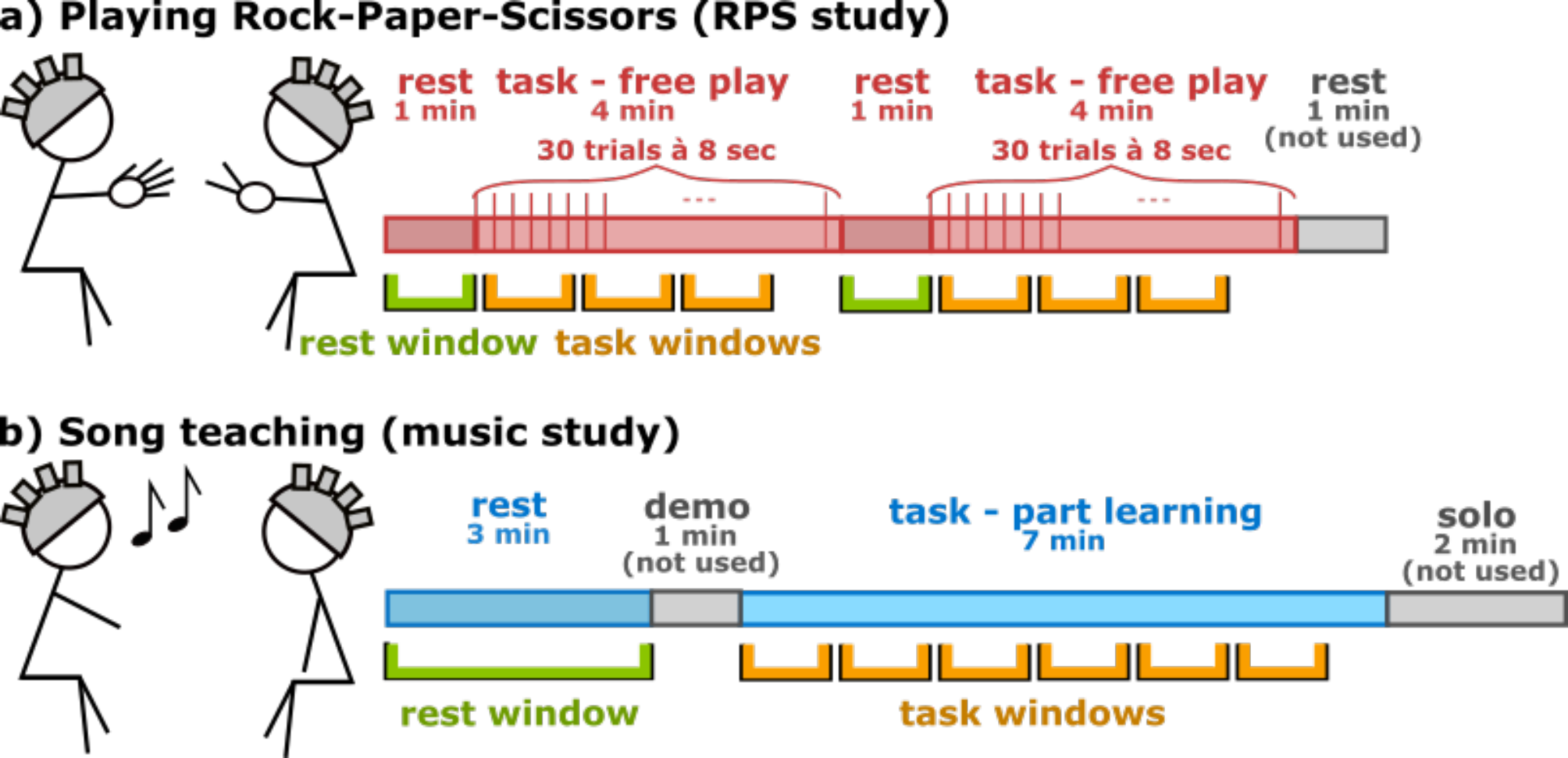
Experimental designs of the data sets and window selection for data re-analysis. a) For the RPS study, subjects played Rock-Paper-Scissors with a timed initiation of trial every 8 seconds (free play condition, see Kayhan et al., 2022) in two blocks of a rest phase followed by a task phase. For the data re-analysis, we used both experimental blocks with the complete rest phase as baseline (green window) and non-overlapping windows of predefined varying sizes (orange windows) for the task phase. b) For the music study, a teacher taught a song to a student partwise (part learning condition, see Pan et al., 2018). For the data re-analysis, we used the complete rest phase as a baseline (green window) and non-overlapping windows of predefined varying sizes (orange windows) for the task phase.

For a near real-time experiment, it is essential to determine the length of the window of integration, meaning how long data needs to be integrated until it can be fed back. The choice of the period of interest directly determines the minimal length of the integration window since the choice of wavelet determines the resolution in frequency and time. Wavelet transformation has a limited resolution in time and frequency/period, which leads to boundary effects at places where the wavelet is longer than the available signal time course. This leads to areas within and outside a “cone of influence” (COI). Within the COI, boundary effects are negligible. For real-time analysis, to avoid boundary effects at the measurement borders, we must discard coherence values outside the COI, since we only have data until the current time point. For a period of 14s, this results in a COI of √2 ∗ 14𝑠 = 19.8𝑠 using the COI definition of Matlab’s wavelet toolbox for Morlet wavelets, which follows the definition of Torrence and Compo (1998). The COI is not only relevant at the measurement boundaries but generally describes the temporal resolution of the wavelet of the corresponding period band. Also, to ensure that no signal from out of our window of interest is leaking in and that all effects measured are only happening within this window, we need to discard about 20 seconds at the beginning and end of our window after determining our coherence. This leads to our choice of 50 seconds as the minimal window investigated. Figure 2c depicts the COI for a time window and the resulting window. Since we want to weigh all periods equally, the size of the resulting window is determined by the COI value for the greatest period in the period band of interest.

**Fig. 2:**
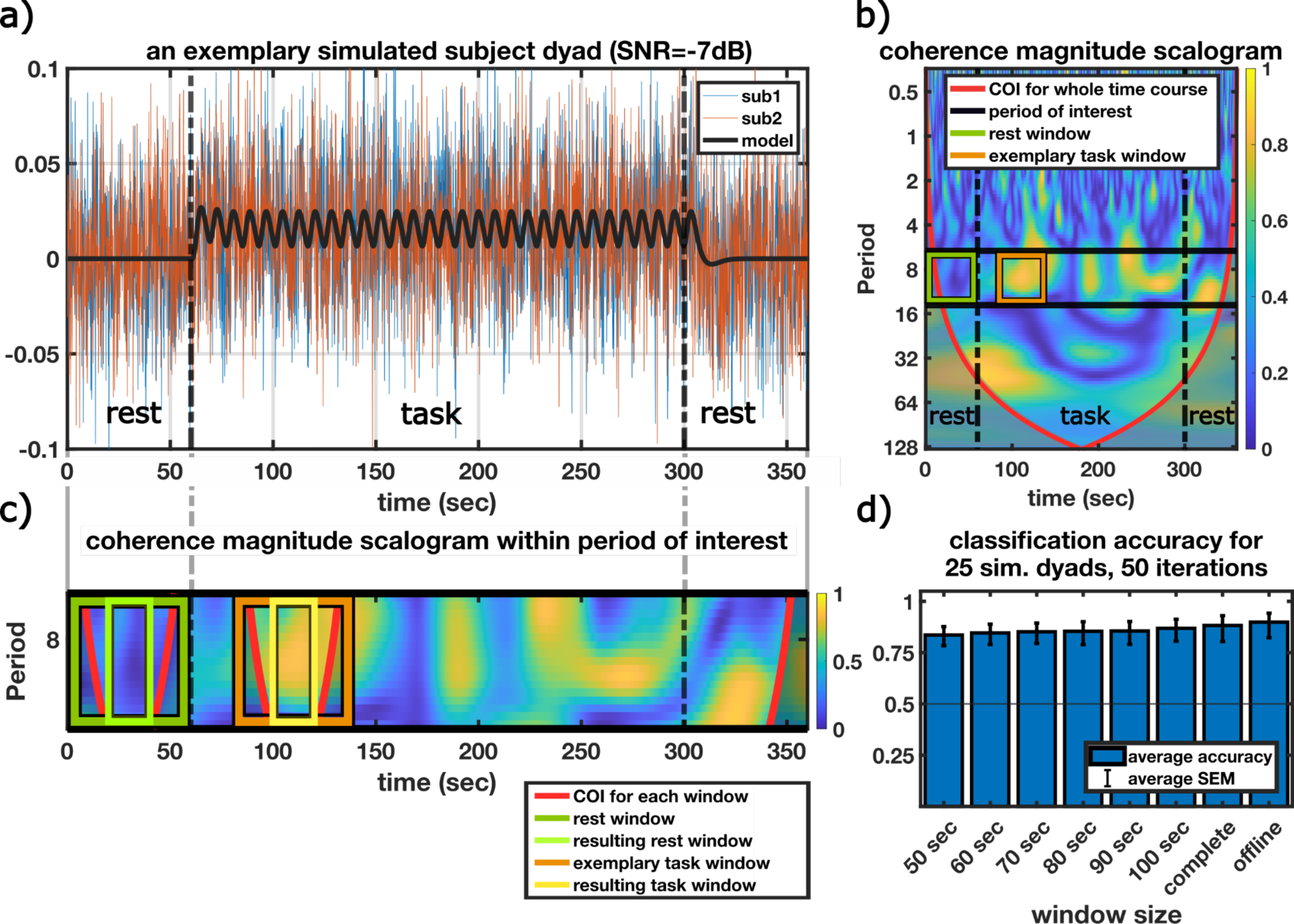
Simulations of subject dyads for a high noise level (SNR = -7 dB). a) An exemplary simulated dyad with a model of zero activity for the rest phase and event-related activity for the task phase plus a high level of added Gaussian noise. b) coherence magnitude scalogram for the dyad depicted in a). The analysis windows are marked in green (rest phase window) and orange (exemplary task window). c) Close-up of the magnitude scalogram of b) for the period band of interest (black lines). The relation between a window and the COI is shown: for the exemplary task window (orange) the COI is plotted (black) and the resulting smaller window (yellow). The same is plotted for the rest block (dark green and light green for the resulting window). d) Classification accuracy of correctly (WTCtask > WTCrest) classified windows for the 25 simulated dyads, rerun for 50 iterations. The bar depicts the average intercept of the mixed-effects model over these 50 iterations, the error bars are the standard errors of the mean of the intercept averaged over 50 iterations. All accuracies reach an average significance level of p<0.001.

### 2.2 Near real-time analyses

A major prerequisite to a neurofeedback study is to assure that the metric chosen for feedback can be obtained in near real-time. In conventional fNIRS studies, the complete time course is wavelet transformed, which is impossible in a real-time setting where data are only available up to the current time point. To evaluate near real-time feasibility, we determined our metric up to the current time point and compared the results for different time windows over which the data was integrated. This was done in simulated data, as well as in existing data where subjects showed higher WTC in the task compared to the rest phases using conventional offline analysis (Kayhan et al., 2022; Pan et al., 2018). We examined if we could reproduce this offline result using a near real-time WTC calculation based on shorter time windows.

As we would do in a near real-time neurofeedback experiment, we took the average WTC values of the complete rest phase right before the task phase as the baseline. For the task phase, we took non-overlapping windows of varying lengths and averaged the WTC values within them. We used an offset of 8 seconds at the start of the task phase and between the windows. The offset accounts for delays due to the hemodynamic response and ensures the independence of the windows of integration. Note that this means that the placement of the windows differed for each window length and that not all data were used for each window analysis. We compared window lengths of 50, 60, 70, 80, 90, and 100 seconds. Due to our choice of the period band of interest and the corresponding COI, 50 seconds is the lower boundary to the length of the window. We did not investigate window durations longer than 100 seconds since such durations would cause a relatively long delay for neurofeedback training. We compared this window of integration analysis with (1) the analysis of a complete block, meaning the rest phase and the task phase together (see Fig. 1), and (2) the whole time course of the entire experiment as it would be analyzed for offline analyses. This “offline” analysis is equivalent to the conventional analysis, meaning the complete time course of the whole experiment was first wavelet transformed, and analyses were performed thereafter.

To assess the success of these analyses, we assumed that the classification and therefore the feedback would be correct if the WTC values were higher during the current task window than during the rest block. To follow this end, we expanded the approach of Koush et al. (2013). Per dyad, we coded every comparison of *WTCtask_window > WTCrest* as a success or failure respectively. We then fitted a generalized linear mixed-effects model to the data using Matlab’s fitglme function. We fitted the formula *success ∼ 1 + (1|dyadID),* with dyadID being the identifier for each dyad to account for the data hierarchy. For the model fit we assumed a binomial distribution using a logit link and used the default maximum pseudo-likelihood as an estimation method. For parts of the music data set, every comparison *WTCtask_window > WTCrest* was a success. To handle such cases, we implemented a continuity correction by adding pseudocounts to reach the solvability of the model estimation (Sweeting et al., 2004). We added both one success and one failure to an arbitrary dyad. In the case of the simulations (see below), also occasionally a near-perfect set, meaning almost every comparison being a success, led to problems with all fitting methods available in Matlab’s fitglme function (i.e. convergence was not possible, which led to unrealistically large p-values). In these cases, we have removed these results from further analysis.

The overall classification was successful if the intercept of the estimated model was significantly different from 0. In the illustrations, we transformed the estimate of the intercept via the inverse logit function to depict the overall classification accuracy/ detectability on a percentage scale.

### 2.3 Simulations

To gain an understanding of the theoretical behavior of the data, we ran simulations that mimicked the design of one of the data sets (Fig. 1a, Kayhan et al., 2022). The design we modeled consisted of a 60-second rest phase, followed by a task phase of 30 events with an intertrial interval of 8 seconds, followed by another 60-second rest phase afterward. This model was convolved with SPM12’s hemodynamic response function (https://www.fil.ion.ucl.ac.uk/spm/software/spm12/). To investigate the effect of different signal-to-noise ratios (SNR) on the results, we simulated 25 subject dyads and added Gaussian noise of a medium SNR of 0 dB and of a high SNR of -7 dB to the convolved model time course for each subject. We then computed the WTC for the different window lengths of 50, 60, 70, 80, 90, and 100 seconds, as well as for the complete block and as in a conventional offline analysis. Afterwards, we computed the WTC classification accuracy (task window > rest). To account for the variability between different sets of 25 dyads of simulated subjects, we reran this analysis 50 times (i.e. performed 50 virtual experiments) to achieve an average success rate and standard error.

### 2.4 Re-analysis of existing data

To assess the performance of real data, we re-analyzed the original data from two existing hyperscanning studies in a pseudo real-time fashion (Fig. 1). The original data collection was carried out under the ethics approval of the Medical Faculty of the Leipzig University (RPS study) and East China Normal University (music study). The original ethics applications included consent for secondary data analysis, as carried out here.

One of the studies, the RPS study, assessed the synchrony between two participants playing Rock-Paper-Scissors (Kayhan et al., 2022). In the study, dyads had to play in different conditions where they either had to predict each other’s actions or play freely. For the analysis in this manuscript, we only used the free play condition as it resulted in the biggest IBS contrast to the resting phase in the original analysis. Here, the dyads had to play rock-paper-scissors as they normally do receiving a prompt to do so every 8 seconds. During this condition, a rest phase of 60 seconds followed a task phase of 4 minutes twice (see Fig. 1a). For this analysis, data from 27 dyads was used.

The second study, the music study, investigated the IBS between teacher and learner while learning a song (Pan et al., 2018). Here one half of the dyads of teacher and learner were to learn a song as a whole, and the other half was to learn a song part-wise, following musical phrases. For this analysis, we only used the “part-learning” condition, as it showed partly higher IBS compared to the rest phase than the other condition, as well as a correlation with subsequent song performance. For this analysis, we used a rest phase of 3 minutes and a task phase of 7 minutes (see Fig. 1b). This analysis is performed on the data of 12 dyads of the part-learning condition.

#### Preprocessing

To increase comparability, we preprocessed both data sets the same way, following the procedure of Kayhan et al. (2022). Specifically, we used the functions of the Homer 2 toolbox (https://homer-fnirs.org/) integrated into Matlab scripts. We first removed channels with nonphysical (negative) intensities. We then corrected the raw optical densities for motion using Homer 2’s motion artifact detection and subsequent cubic spline interpolation (Scholkmann et al., 2010), applied a Butterworth bandpass filter of 3rd order with cutoff values at 0.01 and 0.5 Hz, and converted the data to HbO and HbR values using the modified Lambert-Beer law.

#### Pseudo real-time analysis

For both data sets, we selected the conditions with the biggest difference in WTC reported between the rest block and the task block. We therefore used the “free play” condition of the RPS data set, and the “part learning” condition of the music data set. We then selected the channels that showed a significant effect in the original analysis. In this manuscript, we present the results based on the chromophore used in the original analysis, i.e. HbO for the music study and HbR for the RPS study. The results of the other chromophore can be found in the Supplementary Information (Figures S2 and S3).

Our goal was to investigate whether the difference between task and rest also holds for smaller time windows, as would be used for feedback in a near real-time experiment. We, therefore, re-analyzed the selected data in a pseudo real-time fashion. This approach involves utilizing the data up until the current time point to calculate the WTC windows of integration, meaning wavelet transforming each window of integration. For both data sets, we compared the WTC of the rest block with task block window lengths of 50, 60, 70, 80, 90, and 100 seconds. Additionally, we compared taking the complete task phase as one window of integration and finally the conventional offline analysis. We then determined the classification accuracy as described above.

## 3. Results

### 3.1 Simulations

To assess the theoretical behavior of WTC data for shorter time windows, we simulated artificial subject dyads using an event-related design. Figure 2 depicts the results for our model with a high noise level (SNR=-7dB). It shows a simulated subject dyad (Fig. 2a), the resulting magnitude scalogram for this particular dyad, i.e. the coherence distribution in period and time (Fig. 2b and c), as well as the overall simulation ratios of WTC during the task vs. the rest phase (Fig. 2d). The correspondent results for the model with a moderate noise level (SNR = 0 dB) can be found in Supplementary Figure S1. For no noise (not shown), every window would be correctly classified (WTCtask > WTCrest) regardless of its duration. For a moderate noise level of SNR = 0 dB, i.e. noise and simulated signal both having the same power, we still achieved an almost perfect detection rate (Supplementary Figure S1). Only for a high noise level, the classification rate dropped (Fig. 2d). On average over 50 iterations, meaning 50 virtual experiments with 25 dyads each, we observed a slight decline of correctly classified windows towards the smaller window sizes.

Overall, based on these simulations, windowed WTC for neurofeedback depicted task vs. rest differences reasonably well, and the larger the window length, the better the offline analysis results were approximated even at high noise levels.

### 3.2 Real data re-analysis

To complement the simulations, the same analysis was carried out using the actual data of two interaction studies. Both data sets were analyzed conventionally (“offline” condition) and for shorter windows of integration, as would be the case for neurofeedback. As Fig. 3 shows, overall, the classification accuracy (WTCtask > WTCrest) varied and tended to be lower for the windowed analysis than in the offline condition. However, for all the channels where the offline contrast was significant in our analysis, i.e. the intercept of our success mixed-effects model was significantly different from 0, this result also held for almost all windows tested (Fig. 3 and Supplementary Table 1 and 2). Only channel 16 in the RPS data set did not show significantly higher synchrony for task than rest in our analysis, contrary to the results obtained in the original publication. Note that we did not perform the exact same analysis since in contrast to the original analysis, we treated every channel separately here. In contrast to the simulations, the effect of the window size was less systematic, although for some channels (e.g. channel 10 of the music data set), longer windows steadily increased classification performance. Overall, the near real-time windowed analysis of existing data revealed that for most channels, even shorter windows showed a significantly higher synchrony for task than rest, although the classification performance varied between channels and window sizes.

**Fig. 3:**
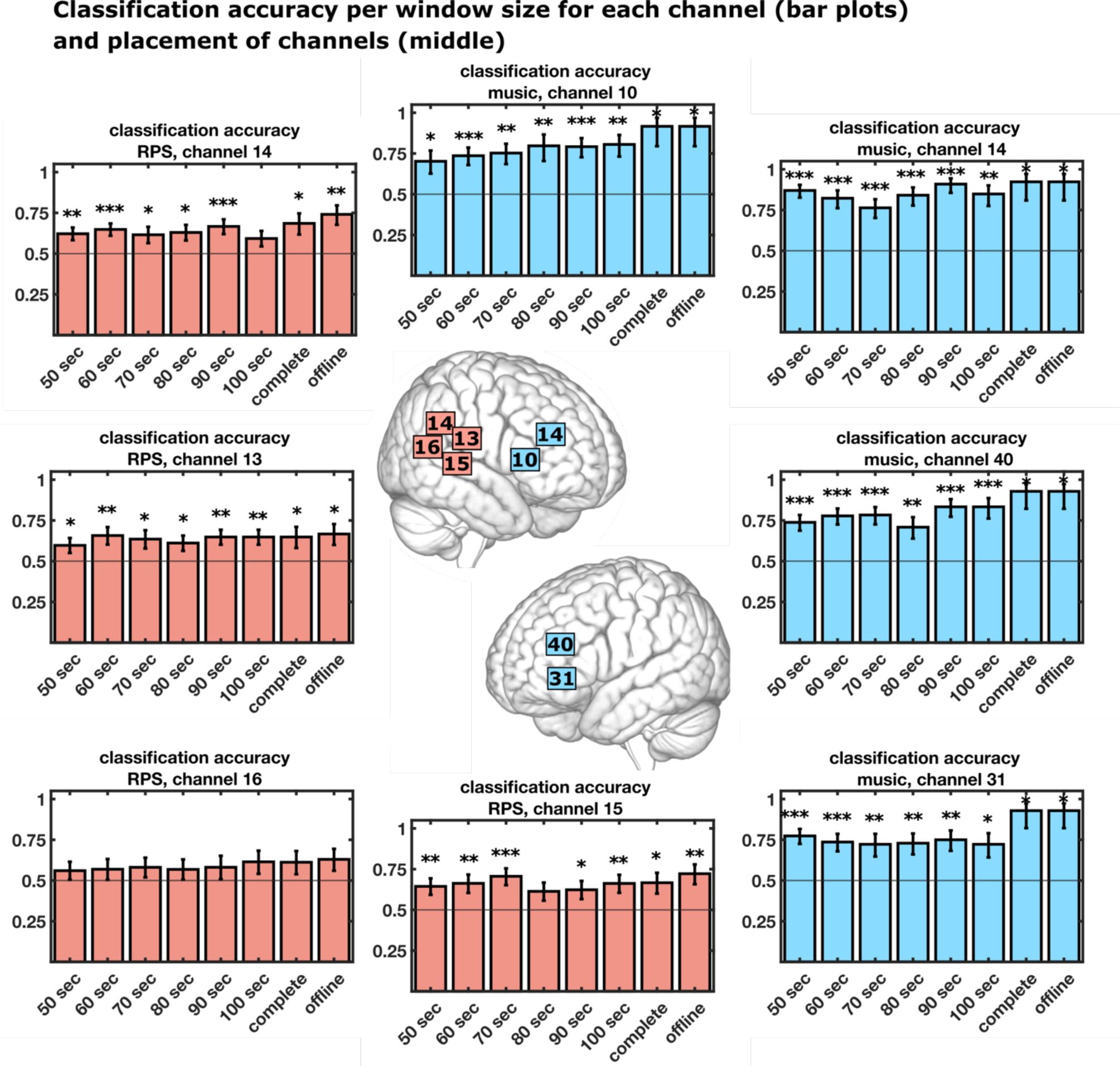
Classification accuracy (percentage of successes) per window size for each channel (bar plots) for the Rock-Paper-Scissors experiment (red) and the music experiment (blue). Most windows are correctly classified (WTCtask > WTCrest). In the middle of the figure, the placement of the channels is depicted. For the Rock-Paper-Scissors experiment, channels within the right temporo-parietal junction, and for the music experiment, channels within the right and left dorsolateral prefrontal cortex and inferior frontal cortex were investigated. The durations on the x-axis indicate the different window sizes ranging from 50 seconds to the complete block as one window. “Offline” means conventionally analyzed. The bar plots show the intercept of the mixed model, the error bars are the standard error of the mean of the intercept. Significance levels: p<0.05*, p<0.01**, p<0.001***. No asterisk means not significant

## 4. Discussion

The aim of this study was to assess the feasibility towards using neurofeedback to train control of interbrain synchrony to improve human interaction. To achieve this goal, we investigated whether the current approach of using WTC in fNIRS setups for investigating interbrain synchrony could be technically adapted for near real-time analysis. Specifically, we examined the feasibility of using WTC for near real-time analysis in fNIRS settings by simulating artificial data and by re-analyzing existing data from two interbrain synchrony experiments.

### 4.1 Feasibility

Our results show that for the selected channels with significantly higher WTC, and thus IBS, in the conventional offline analysis, the WTC is also higher in the windowed analysis. This means that using the WTC signal obtained from shorter time windows, as would be the case in near real-time analysis and necessary for neurofeedback, seems technically feasible and stable enough. For both the simulations and the re-analysis of existing data, the detectability of higher WTC in the task condition was lower for the near real-time analysis. This was expected, since there are fewer data points available for analysis, and noise effects, therefore, become more visible. Indeed, the simulations show that increased window length and reduced noise levels increase performance (Fig. 2 and S1). For the re-analysis of existing data, the effect of window length is less pronounced. This might be because the variability is generally high due to varying noise levels in experimental settings. Also, in our simulations, in the high noise condition, the variability between different iterations is relatively high. Therefore, it can happen that for one iteration of 25 dyads, a shorter window size actually yields a higher contrast than a longer window size. Overall, our results of the real-data re-analysis are consistent with our simulations in the high-noise condition.

These results suggest that shorter window durations likely contain meaningful information for fNIRS hyperfeedback and longer window durations of up to 100 seconds provide limited benefits. Longer integration durations can be problematic for neurofeedback, because the number of feedback presentations is reduced and the temporal contingency between the participants’ actions and the feedback is important for learning self-regulation. Therefore, our results suggest that for our period band chosen, which is a classical range in fNIRS IBS studies, a window size of around one minute seems advisable.

Given these results, we recommend intermittent neurofeedback instead of continuous neurofeedback. This form of intermittent feedback has been successfully used in the past and in some settings even worked better than continuous feedback (Johnson et al., 2012; Watanabe et al., 2017; Hellrung et al., 2018; but see also Emmert et al., 2017). The advantages of intermittent feedback are that more time is available for complex feedback signal computations and feedback accuracy/validity is improved due to averaging over more data points. Importantly, the participants can focus on self-regulation without the simultaneous processing of a feedback signal, which is known to cause dual-task interference.

We have chosen to operationalize IBS by determining WTC since it is by far the most used metric in fNIRS hyperscanning (Hakim et al., 2023). However, generally, no agreement on measuring IBS has been reached yet (Andrea et al., 2022; Ayrolles et al., 2021; Hakim et al., 2023; Holroyd, 2022) and caution should be taken when generalizing this result to other IBS metrics. Classical correlation measures and WTC do not yield the exact same results in all cases. The Pearson correlation coefficient is dependent on the exact relation of the phases of two signals, whereas WTC does not depend on the exact phase relation as long as the signal is phase-locked, meaning that the phase relation does not vary over time. This makes exact comparisons of synchrony results assessed with different metrics difficult. It also implies a slightly different underlying definition of synchrony: with WTC, two signals in one period band can have a constant phase shift up to the corresponding period, and one signal can lead or follow the other. Synchrony will be high as long as this phase difference between the two signals remains constant; see also Hakim et al. (2023) for a recent review of IBS metrics. Our results, therefore, connect to a substantial body of literature that applied WTC, but cannot be automatically generalized to other synchrony metrics.

The choice of WTC as a metric for IBS also influences the length of the window of integration. Since a signal cannot be localized equally well in frequency/period and time (due to the physical Heisenberg-Gabor limit, Gabor (1946)), our lowest period band of interest determines the temporal resolution. The higher the period, the wider the wavelet in use and the wider the temporal integration of the signal for that period by the signal processing algorithm. This is the wavelet’s COI, which generally describes the “resolution” of a wavelet and therefore the temporal resolution of the derived coherence values. This is especially relevant for non-periodic signals, as usually is the case with fNIRS data. Note that the temporal resolution could theoretically be improved by worsening the period resolution. This would mean deviating from the WTC analysis as it is currently used in the field and is therefore beyond the scope of this manuscript.

For this study, we can conclude that the WTC metric as it is currently practiced in the field can technically be reasonably well assessed using windows of integration that would be suitable for intermittent neurofeedback. However, whether subjects can learn to self-regulate and learn from such a neurofeedback signal also depends on other experimental factors. Such factors would be the feedback modality, the clarity of instructions, the training duration, as well as subject-specific factors such as individual motivation, height of baseline IBS, and effective regulation strategies. How subjects learn to regulate their IBS is an important topic for future research towards which the assessment of technical feasibility is an essential prerequisite.

### 4.2 Limitations

One limitation of this study is the preprocessing of the data, i.e. the motion correction, filtering, and transformation of raw signal to HbO/HbR values. For the analyses in this manuscript, the data was preprocessed in a conventional offline way, which will need to be adapted for a hyperfeedback experiment. All preprocessing steps will need to be performed online in near real-time. This means that, in particular, the bandpass filter used here needs to be exchanged by a causal filter. Also, artifact control is essential and should be improved, e.g., by utilizing short-distance channels to measure physiological artifacts online and filter them out (Godet et al., 2023; Klein et al., 2022). Note that there are already different approaches for real-time preprocessing in practice (Kohl et al. 2020).

Another limitation is that the channels that were re-analyzed here were those that produced significant task effects in the original offline analyses. Such a pre-selection of channels would not be possible in a hyperfeedback setting. However, in neurofeedback studies the brain target definition is often based on pre-training localizer runs, pilot studies, or existing literature (Sitaram et al., 2017; Sulzer et al., 2013). Similarly, in hyperfeedback, the brain target can be defined a *priori* using such approaches to brain target definition.

Finally, in our simulations, we have only assumed Gaussian noise. In the future, more specifically tailored noise models could be used to achieve more exact simulations (Gemignani and Gervain, 2021). Additionally, the model assumed for the simulations is a very simple one only modeling each task event onset, likely discarding additionally ongoing activity in the brain areas investigated, which are often discussed in processing social tasks. The simulations should, therefore, be taken as a first proof of concept that a near real-time analysis is feasible, and less as an exact model of the RPS experiment.

### 4.3 Conclusion

Our feasibility study provides evidence that a near real-time fNIRS hyperfeedback setup for IBS, measured in terms of WTC, is feasible. Given the relatively long windows needed for this analysis, an intermittent neurofeedback setup seems most suitable. In the future, it will be important to improve the setup and processing pipeline to remove noise as much as possible. Once the technical challenges have been overcome, it will be important to investigate if and how subjects can make good use of this new type of neurofeedback, to realize the full potential of hyperfeedback.

## Code and data availability statement

All code used for this manuscript is openly available. The code for preprocessing the music data and all further analyses for this manuscript can be found at https://github.com/univiemops/hyperfeedback-feasibility

The code for the preprocessing of the RPS data set can be found at https://github.com/tnguyen1992/RPS

The data re-analyzed for this secondary analysis were obtained from Pan et al. (2018) and Kayhan et al. (2022), and the data availability of the original publications applies.

## Acknowledgments

The authors would like to thank Ezgi Kayhan for providing the data of the RPS study.

KK is funded by the Austrian Science Fund (FWF) ESPRIT Programme (ESP 289). TN is funded by the European Union (MSCA, SYNCON,101105726). FM is funded by the Austrian Science Fund (FWF) ESPRIT Programme (ESP 133). BS is funded by The Netherlands Organization for Scientific Research (NWO Vidi-Grant VI.Vidi.191.210).

## CRediT author statement

Conceptualization: KK, FS

Methodology: KK, TN, DS, FM, FS

Software: KK, TN

Resources: YP, YH, TN, SH, FS

Data curation: TN, SH, YP, YH

Formal Analysis: KK Visualization: KK

Supervision: BS, SH, FS

Writing – Original draft: KK

Writing – Review and editing: KK, TN, DS, FM, YP, BS, SH, FS

