## Supplementary Material for "Towards fNIRS Hyperfeedback: A Feasibility Study on Real-Time Interbrain Synchrony"

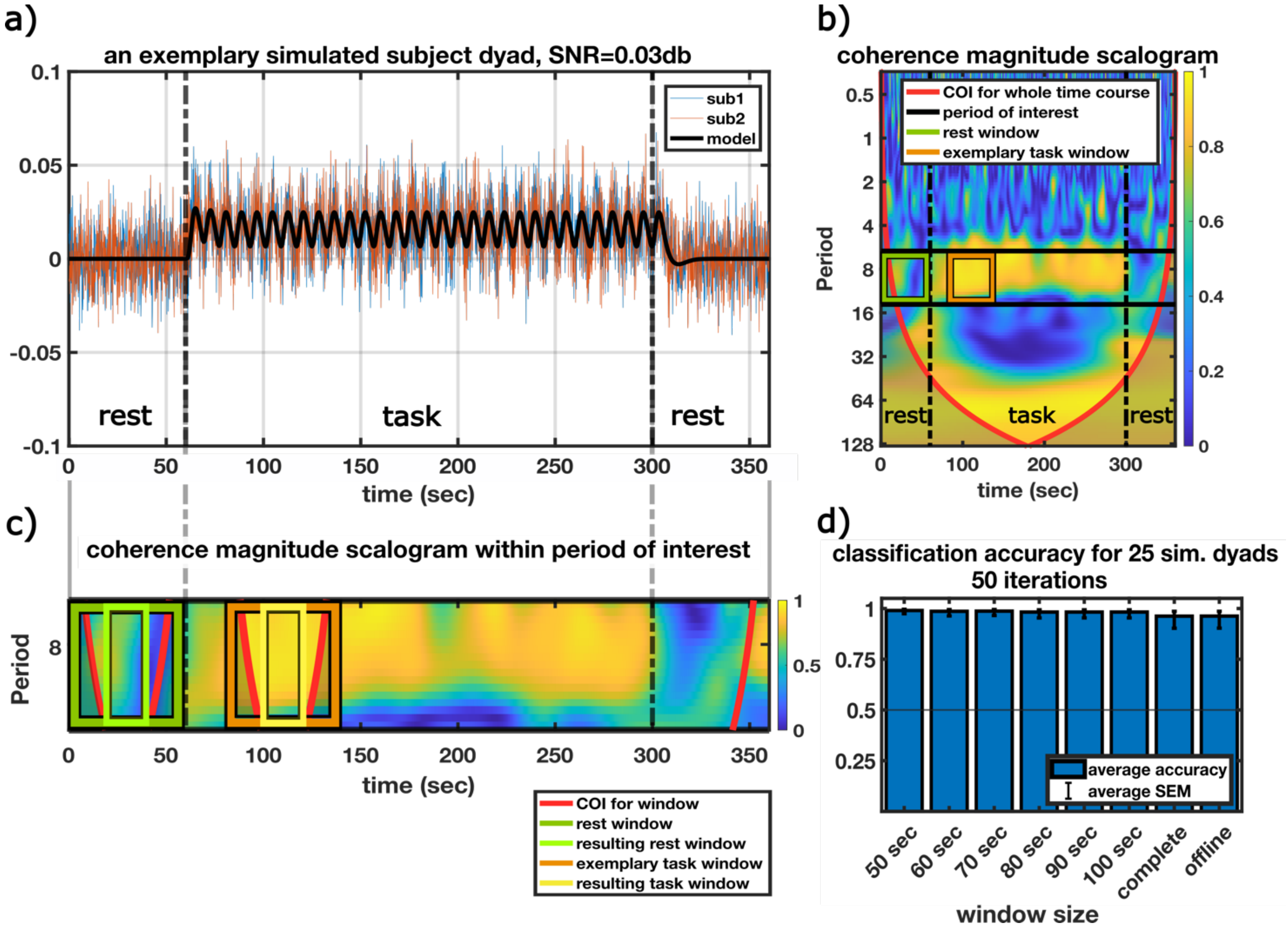


Supplementary Figure S1: Simulated data with a moderate noise level. Figure analog to Fig. 2.

| music - channel 10 - HbO | | | | | | | | |
|---|---|---|---|---|---|---|---|---|
| **Row** | **windowsize 50 sec** | **windowsize 60 sec** | **windowsize 70 sec** | **windowsize 80 sec** | **windowsize 90 sec** | **windowsize 100 sec** | **complete block** | **offline** |
| **estimate** | 0.857251747 | 1.026686977 | 1.11205597 | 1.367353062 | 1.335001067 | 1.421385681 | 2.397895273 | 2.397895273 |
| **SE** | 0.339531562 | 0.27622823 | 0.336211844 | 0.501968717 | 0.355409327 | 0.421117444 | 1.044465942 | 1.044465942 |
| **tStat** | 2.524807243 | 3.716806856 | 3.307604979 | 2.723980631 | 3.756235319 | 3.375271435 | 2.295809923 | 2.295809923 |
| **DF** | 83 | 71 | 59 | 47 | 47 | 35 | 11 | 11 |
| **pValues** | 0.013479606 | 0.000399246 | 0.001606185 | 0.009026829 | 0.000475189 | 0.001816756 | 0.042342867 | 0.042342867 |
| **Lower** | 0.181937126 | 0.475903531 | 0.439297624 | 0.357522258 | 0.620009725 | 0.566471818 | 0.099041235 | 0.099041235 |
| **Upper** | 1.532566368 | 1.577470423 | 1.784814316 | 2.377183866 | 2.049992409 | 2.276299544 | 4.696749311 | 4.696749311 |

| music - channel 14 - HbO | | | | | | | | |
|---|---|---|---|---|---|---|---|---|
| **Row** | **windowsize 50 sec** | **windowsize 60 sec** | **windowsize 70 sec** | **windowsize 80 sec** | **windowsize 90 sec** | **windowsize 100 sec** | **complete block** | **offline** |
| **estimate** | 1.902368544 | 1.533468513 | 1.172720261 | 1.665007764 | 2.302585093 | 1.722766598 | 2.48490665 | 2.48490665 |
| **SE** | 0.343047272 | 0.371941815 | 0.317384004 | 0.412170075 | 0.524404424 | 0.485504163 | 1.040833 | 1.040833 |
| **tStat** | 5.545499699 | 4.122872047 | 3.694957047 | 4.039613417 | 4.390857489 | 3.548407467 | 2.387421085 | 2.387421085 |
| **DF** | 76 | 65 | 54 | 43 | 43 | 32 | 12 | 12 |
| **pValues** | 4.07437E-07 | 0.000108278 | 0.000514678 | 0.000216937 | 7.22783E-05 | 0.001221452 | 0.034296058 | 0.034296058 |
| **Lower** | 1.219130688 | 0.790649414 | 0.536403646 | 0.833787589 | 1.245022781 | 0.733826979 | 0.217126356 | 0.217126356 |
| **Upper** | 2.585606399 | 2.276287612 | 1.809036876 | 2.496227938 | 3.360147405 | 2.711706216 | 4.752686944 | 4.752686944 |

| music - channel 31 - HbO | | | | | | | | |
|---|---|---|---|---|---|---|---|---|
| **Row** | **windowsize 50 sec** | **windowsize 60 sec** | **windowsize 70 sec** | **windowsize 80 sec** | **windowsize 90 sec** | **windowsize 100 sec** | **complete block** | **offline** |
| **estimate** | 1.229948291 | 1.026725244 | 0.95538559 | 0.990398704 | 1.098612289 | 0.955511445 | 2.56494936 | 2.56494936 |
| **SE** | 0.260799149 | 0.27640715 | 0.347096246 | 0.324799178 | 0.333333335 | 0.372104205 | 1.037749044 | 1.037749044 |
| **tStat** | 4.716074794 | 3.714539384 | 2.752509139 | 3.049264815 | 3.295836845 | 2.567859841 | 2.471647046 | 2.471647046 |
| **DF** | 83 | 71 | 59 | 47 | 47 | 35 | 13 | 13 |
| **pValues** | 9.59949E-06 | 0.000402251 | 0.007845845 | 0.00376123 | 0.001872239 | 0.014660983 | 0.028047631 | 0.028047631 |
| **Lower** | 0.711229274 | 0.475585041 | 0.260847605 | 0.33698704 | 0.428032113 | 0.200099748 | 0.323028851 | 0.323028851 |
| **Upper** | 1.748667308 | 1.577865447 | 1.649923574 | 1.643810368 | 1.769192464 | 1.710923142 | 4.806869869 | 4.806869869 |

| music - channel 40 - HbO | | | | | | | | |
|---|---|---|---|---|---|---|---|---|
| **Row** | **windowsize 50 sec** | **windowsize 60 sec** | **windowsize 70 sec** | **windowsize 80 sec** | **windowsize 90 sec** | **windowsize 100 sec** | **complete block** | **offline** |
| **estimate** | 1.036091932 | 1.252762968 | 1.285198244 | 0.887303195 | 1.609437912 | 1.609437912 | 2.56494936 | 2.56494936 |
| **SE** | 0.248160389 | 0.283473355 | 0.313368271 | 0.317553676 | 0.387298341 | 0.447213596 | 1.037749044 | 1.037749044 |
| **tStat** | 4.175089895 | 4.41933235 | 4.101239225 | 2.794183356 | 4.155550756 | 3.598812572 | 2.471647046 | 2.471647046 |
| **DF** | 83 | 71 | 59 | 47 | 47 | 35 | 13 | 13 |
| **pValues** | 7.30906E-05 | 3.48171E-05 | 0.000127626 | 0.007507035 | 0.000136065 | 0.000978783 | 0.028047631 | 0.028047631 |
| **Lower** | 0.542510899 | 0.687533152 | 0.658149783 | 0.2484676 | 0.83029415 | 0.701546045 | 0.323028851 | 0.323028851 |
| **Upper** | 1.529672964 | 1.817992785 | 1.912246706 | 1.52613879 | 2.388581675 | 2.51732978 | 4.806869869 | 4.806869869 |

Supplementary Table 1: statistical results of the generalized linear mixed-effects model for the music data set.

| RPS - channel 13 - HbR | | | | | | | | |
|---|---|---|---|---|---|---|---|---|
| Row | windowsize 50 sec | windowsize 60 sec | windowsize 70 sec | windowsize 80 sec | windowsize 90 sec | windowsize 100 sec | complete block | offline |
| estimate | 0.39282174 | 0.65091975 | 0.555516102 | 0.451985124 | 0.610909082 | 0.610909082 | 0.610909082 | 0.693147181 |
| SE | 0.191223476 | 0.237832521 | 0.242624297 | 0.197385509 | 0.20149815 | 0.20149815 | 0.284961416 | 0.288675135 |
| tStat | 2.05425478 | 2.736882861 | 2.289614471 | 2.289859705 | 3.031834701 | 3.031834695 | 2.143830878 | 2.401132266 |
| DF | 215 | 161 | 161 | 107 | 107 | 107 | 53 | 53 |
| pValues | 0.041160595 | 0.006899737 | 0.023342411 | 0.023991301 | 0.003049498 | 0.003049498 | 0.036653077 | 0.019885228 |
| Lower | 0.015908963 | 0.181246163 | 0.076379677 | 0.060691373 | 0.211462501 | 0.211462501 | 0.039348864 | 0.114138185 |
| Upper | 0.769734517 | 1.120593336 | 1.034652528 | 0.843278875 | 1.010355663 | 1.010355664 | 1.182469301 | 1.272156176 |
| **RPS - channel 14 - HbR** | | | | | | | | |
| Row | windowsize 50 sec | windowsize 60 sec | windowsize 70 sec | windowsize 80 sec | windowsize 90 sec | windowsize 100 sec | complete block | offline |
| estimate | 0.497024037 | 0.610909082 | 0.470925547 | 0.531149015 | 0.693147181 | 0.374700361 | 0.778972973 | 1.049822124 |
| SE | 0.16480937 | 0.164522552 | 0.211752969 | 0.205532815 | 0.204124146 | 0.196673242 | 0.301363523 | 0.310529504 |
| tStat | 3.015751083 | 3.713223963 | 2.223938341 | 2.584254075 | 3.395713813 | 1.905192376 | 2.584828336 | 3.380748399 |
| DF | 215 | 161 | 161 | 107 | 107 | 107 | 53 | 53 |
| pValues | 0.00287203 | 0.000281662 | 0.027543963 | 0.011106531 | 0.000961509 | 0.059439503 | 0.012531602 | 0.001363735 |
| Lower | 0.172175024 | 0.286008615 | 0.05275407 | 0.12370418 | 0.288494869 | -0.015181403 | 0.174514294 | 0.426978815 |
| Upper | 0.82187305 | 0.935809549 | 0.889097023 | 0.93859385 | 1.097799492 | 0.764582126 | 1.383431652 | 1.672665434 |
| **RPS - channel 15 - HbR** | | | | | | | | |
| Row | windowsize 50 sec | windowsize 60 sec | windowsize 70 sec | windowsize 80 sec | windowsize 90 sec | windowsize 100 sec | complete block | offline |
| estimate | 0.595138462 | 0.676302227 | 0.876566551 | 0.462808087 | 0.504317252 | 0.673659049 | 0.693147181 | 0.955511445 |
| SE | 0.220152388 | 0.25271751 | 0.24821484 | 0.23438959 | 0.237436605 | 0.24738967 | 0.288675708 | 0.303821815 |
| tStat | 2.703302318 | 2.676119389 | 3.531483247 | 1.974524922 | 2.12400802 | 2.723068627 | 2.401127502 | 3.14497313 |
| DF | 215 | 161 | 161 | 107 | 107 | 107 | 53 | 53 |
| pValues | 0.007414333 | 0.008218386 | 0.000539216 | 0.050898806 | 0.035973394 | 0.007554229 | 0.019885461 | 0.002722283 |
| Lower | 0.161205088 | 0.177233645 | 0.386389875 | -0.001841938 | 0.033626876 | 0.183237881 | 0.114137036 | 0.346122056 |
| Upper | 1.029071835 | 1.17537081 | 1.366743226 | 0.927458112 | 0.975007629 | 1.164080217 | 1.272157325 | 1.564900834 |
| **RPS - channel 16 - HbR** |  |  |  |  |  |  |  |  |
| Row | windowsize 50 sec | windowsize 60 sec | windowsize 70 sec | windowsize 80 sec | windowsize 90 sec | windowsize 100 sec | complete block | offline |
| estimate | 0.243362504 | 0.278719911 | 0.326813202 | 0.274650542 | 0.32773209 | 0.466640658 | 0.456430105 | 0.531227481 |
| SE | 0.22513199 | 0.261796658 | 0.24908416 | 0.254903332 | 0.298755023 | 0.301539623 | 0.30104489 | 0.288625666 |
| tStat | 1.080977005 | 1.064642742 | 1.312059357 | 1.077469409 | 1.096992737 | 1.547526835 | 1.51615297 | 1.840541378 |
| DF | 215 | 161 | 161 | 107 | 107 | 107 | 53 | 53 |
| pValues | 0.280918364 | 0.288631723 | 0.191367741 | 0.283693795 | 0.275107243 | 0.124688951 | 0.135422808 | 0.071290164 |
| Lower | -0.200385958 | -0.238278247 | -0.165080212 | -0.230665583 | -0.264514899 | -0.131126476 | -0.147389479 | -0.047682293 |
| Upper | 0.687110967 | 0.79571807 | 0.818706616 | 0.779966668 | 0.91997908 | 1.064407791 | 1.060249688 | 1.110137255 |

Supplementary Table 2: statistical results of the generalized linear mixed-effects model for the RPS data set.

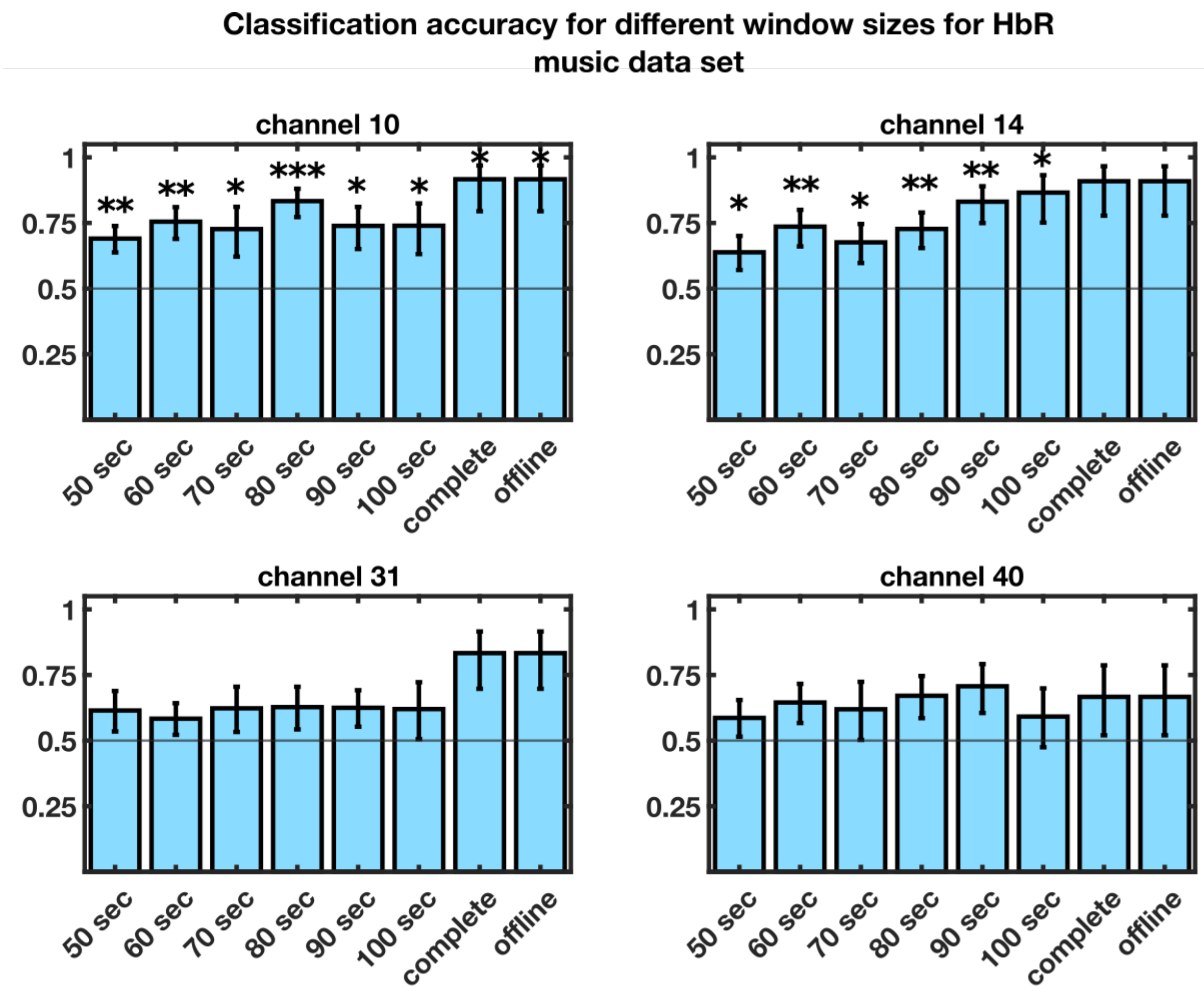


Supplementary Figure S2: Classification accuracy ($WTC_{task}$>$WTC_{rest}$) f for different window sizes for the music/ song teaching data set for HbR. Error bars show the standard error of the fixed effect. Significance levels: p<0.05*, p<0.01**, p<0.001***. No asterisk means not significant.

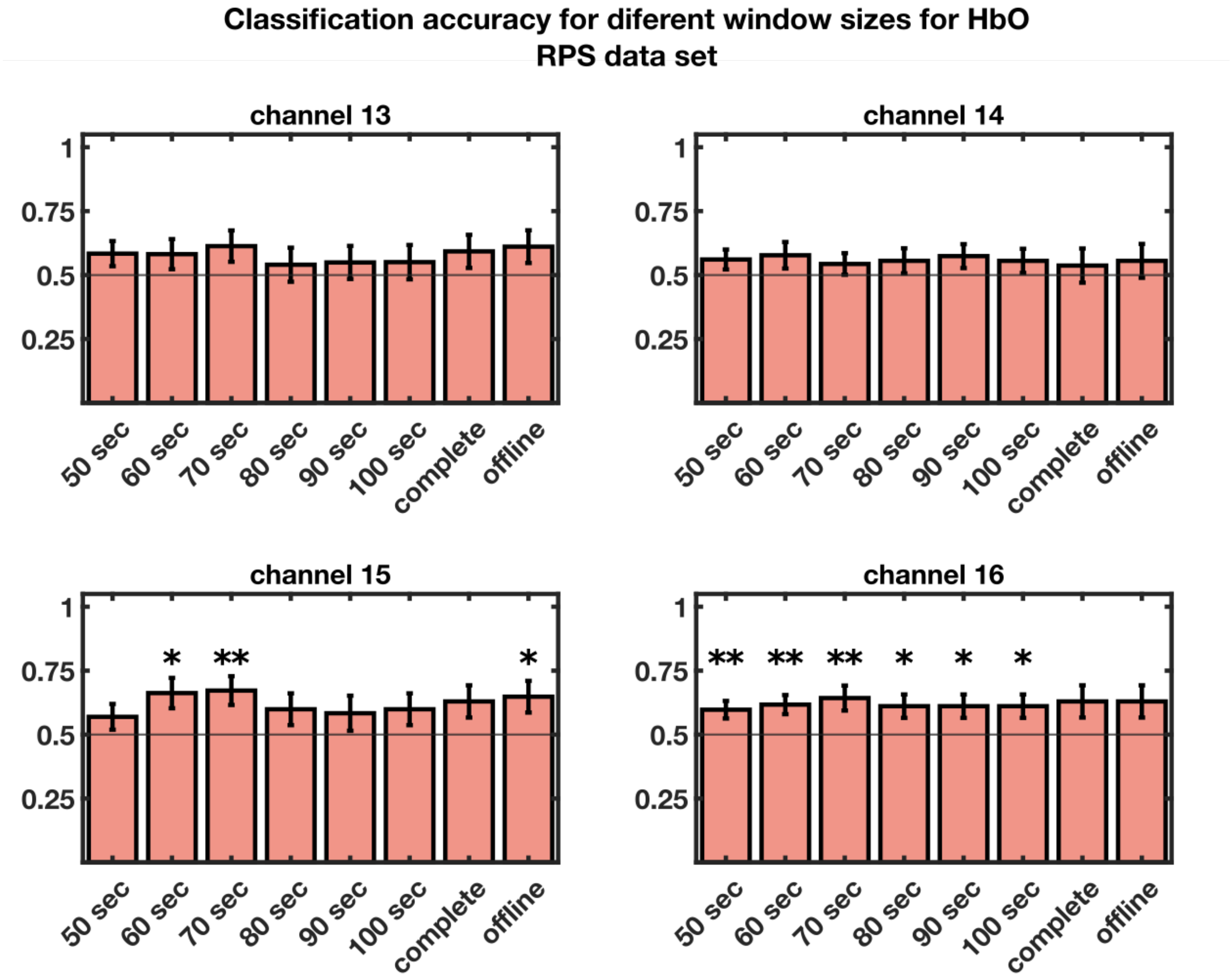


Supplementary Figure S3: Accuracy of classification ($WTC_{task}$>$WTC_{rest}$) for HbO for different window sizes for the RPS data set. Error bars show the standard error of the fixed effect. Significance levels: p<0.5*, p<0.01**, p<0.001***. No asterisk means not significant.

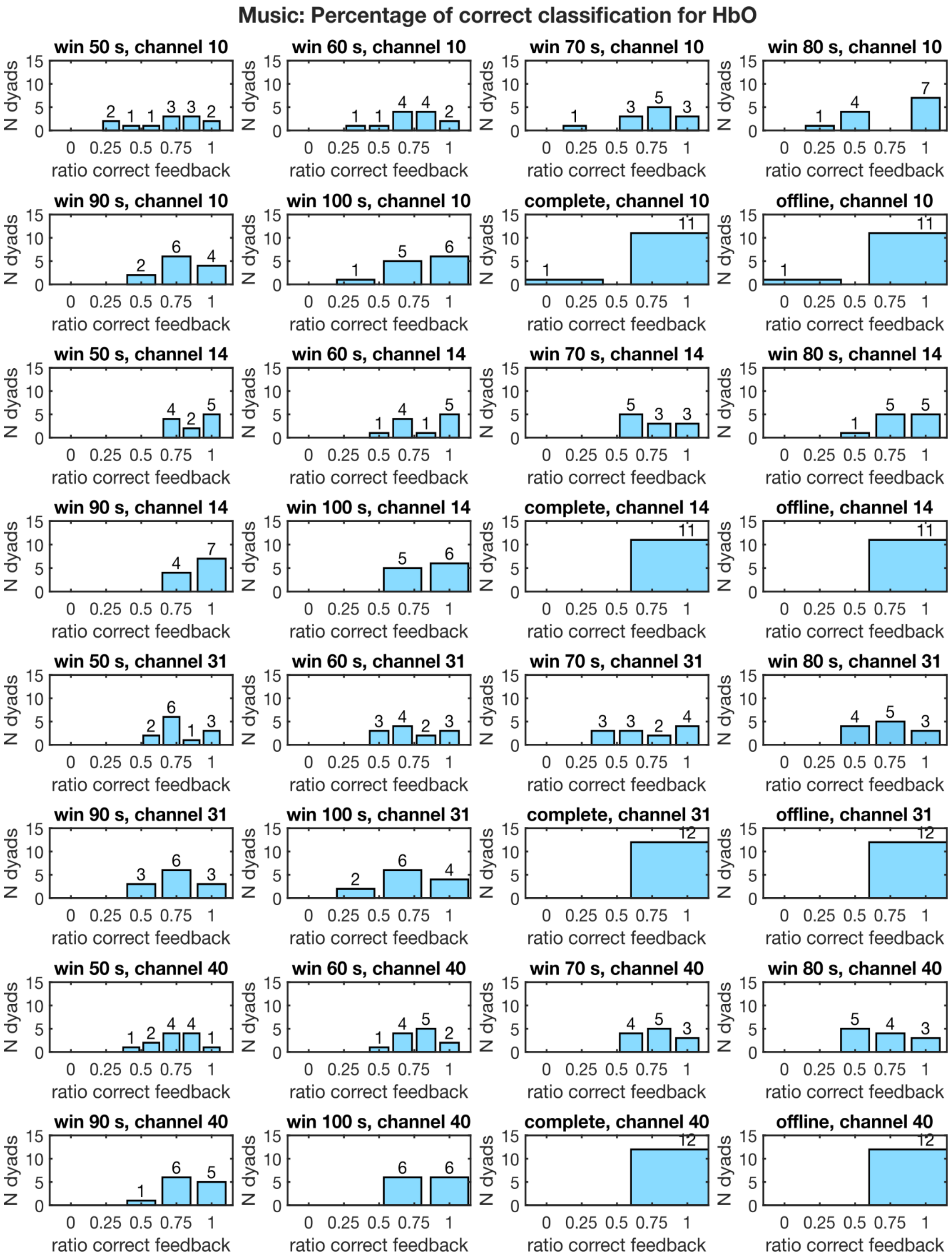


Supplementary Figure S4: Music data set: ratio of correct classification broken down by subject pairs/dyads. Correct classification would mean correct feedback in a neurofeedback experiment. For each window length and channel, it is summarized which individual ratio is achieved for how many dyads. The amount of dyads is also printed on top of each ratio bar. For the first panel, this means: 2 dyads have a ratio of 0.3, 1 dyad a ratio of 0.4, 1 dyad a ratio of 0.6, 3 dyads a ratio of 0.7, 3 dyads a ratio of 0.9, and 2 dyads of 1. 0.5 depicts chance level. Note that due to the total number of windows varying by window length, the possible ratios are also changing.

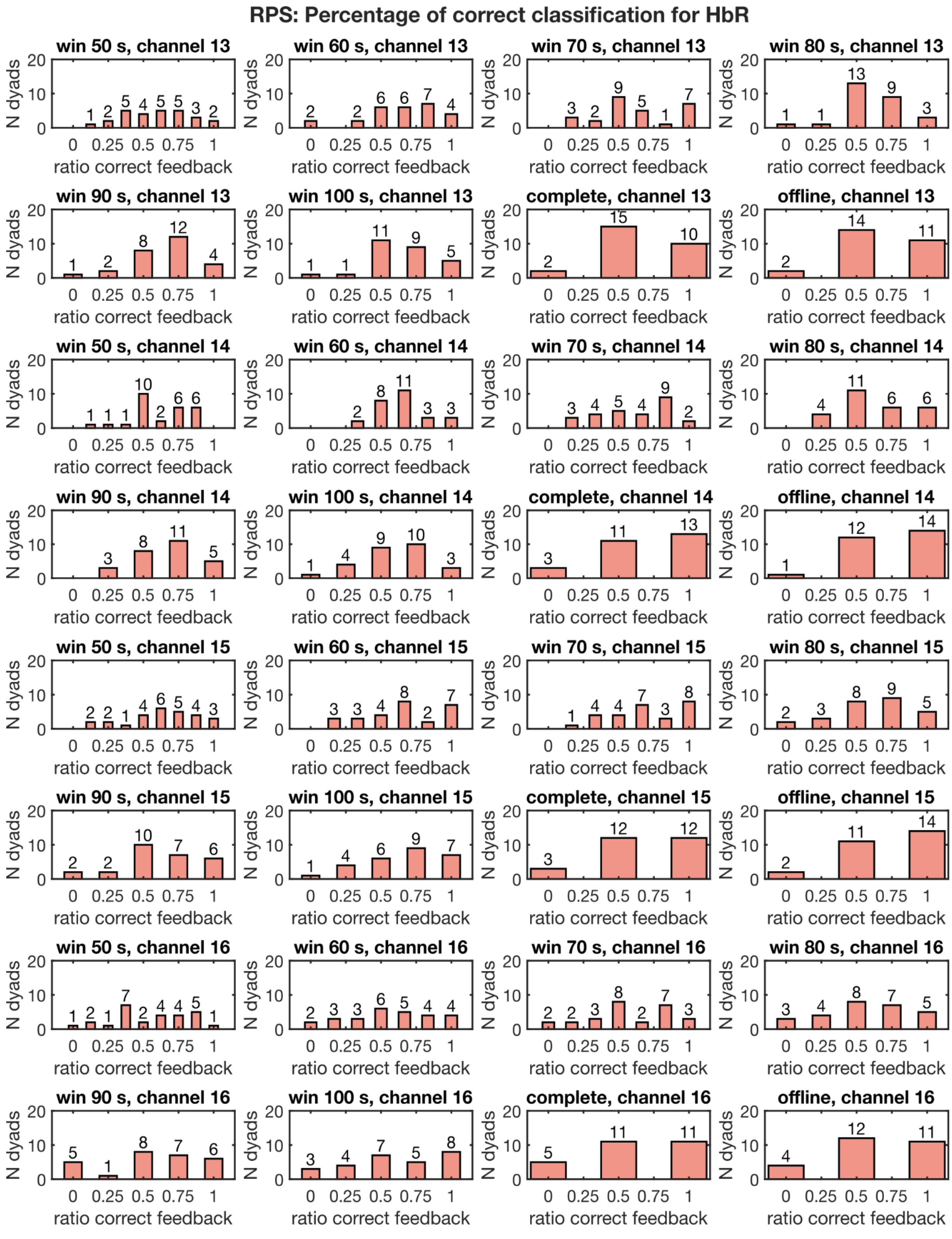


Supplementary Figure S5: RPS data set: ratio of correct classification broken down by subject pairs/dyads. Correct classification would mean correct feedback in a neurofeedback experiment. For each window length and channel, it is summarized which individual ratio is achieved for how many dyads. Note that due to the total number of windows varying by window length, the possible ratios are also changing.
